# Quantum Dot-Based Non-invasive Monitoring of Sinusoidal Endothelial Scavenger Function in Steatohepatitis

**DOI:** 10.64898/2026.11.01.763501

**Authors:** Tomoki Yagai, Muneyo Mio, Sakura Suga, Eri Wada, Hiro Kohda, Kozue Ochi, Ibuki Shirakawa, Xunmei Yuan, Ayaka Ito, Miyako Tanaka, Satoko Arakawa, Yoshinobu Baba, Hiroshi Yukawa, Takayoshi Suganami

## Abstract

Metabolic dysfunction-associated steatotic liver disease (MASLD) and its progressive form, metabolic dysfunction-associated steatohepatitis (MASH), represent urgent medical challenges. Current non-invasive assessments rely on serum indices or tissue stiffness, and none visualizes functional changes within the liver, hindering drug discovery and patient management. In this study, we developed a non-invasive technique using semiconductor quantum dots, as blood-borne foreign particle mimics, to evaluate the scavenger function of liver sinusoidal endothelial cells (LSECs) via near-infrared optical imaging. Progression from MASLD to MASH induced a significant decline in the scavenger capability of LSECs, resulting in a marked reduction of intrahepatic quantum dot retention. Mechanistically, we demonstrated that quantum dots are recognized and internalized by scavenger receptor class A (SR-A) on LSECs, and that SR-A-mediated uptake progressively diminishes with increasing MASH severity. Crucially, quantum dot retention levels significantly correlated with both the progression and regression of MASH pathology. These findings demonstrate that quantum dot-based optical evaluation of LSEC scavenger function offers a highly sensitive, non-invasive diagnostic platform for staging MASH and assessing therapeutic efficacy.

## Introduction

Metabolic dysfunction-associated steatotic liver disease (MASLD), defined as hepatic steatosis with at least one of the cardiometabolic risk factors, such as obesity, type 2 diabetes, hypertension, and dyslipidemia, represents a major global health crisis^1, 2, 3^, affecting approximately 38% of adults worldwide^4^. Among MASLD patients, 12-40% progress to metabolic dysfunction-associated steatohepatitis (MASH), characterized by hepatic inflammation and hepatocellular injury with or without fibrosis^5, 4^. Accurate non-invasive assessment of MASH remains a critical challenge^6, 7^. While vibration-controlled transient elastography measures liver stiffness and lipid accumulation, it lacks the ability to detect inflammation, limiting its ability to distinguish hepatic steatosis from steatohepatitis^8^. Blood-based indices, such as the FIB-4 index, and serum markers, including ALT and AST, likewise lack reliability evaluating MASH severity^9^. Liver biopsy, the gold standard, is invasive and limited by sampling variability and inter-pathologist disagreement^10, 11^. Consequently, surrogate endpoints for drug development and clinical practice remain elusive^12^, and novel strategies capturing functional changes of hepatic resident cells are required.

The hepatic reticuloendothelial system (hRES), consisting of liver sinusoidal endothelial cells (LSECs) and Kupffer cells (KCs), the hepatic resident macrophages, has significant roles in blood clearance, eliminating waste and foreign products^13^. LSECs express multiple scavenger receptors (SRs) that internalize metabolic wastes, damage-associated molecular patterns, and pathogen-associated molecular patterns into lysosomes to degrade them^13, 14^, whereas KCs additionally exhibit vigorous phagocytic capacity^15^. Previous studies unveiled that sinusoidal capillarization in MASH leads to morphological and phenotypic alterations in LSECs, especially a concomitant downregulation of SRs^16^. Although LSEC scavenger function is projected to be important for MASLD and MASH pathogenesis^17^, how it is altered during disease progression remains elusive, because non-invasive method is currently unavailable to measure this function in vivo.

Systemic administration of a contrast agent recognized by SRs offers a potential strategy for functional imaging. Quantum dots (QDs) are a class of quantum sensors composed of semiconductor nanoparticles that exhibit size-tunable fluorescence across a broad spectrum^18^. Although QDs are now implemented in commercial display technologies, their translation into medicine has progressed more slowly, remaining largely at the preclinical stage across regenerative medicine, cancer treatment, and infectious disease research^19, 20, 21, 22–24, 25–27^. In particular, there is no established use of QDs in MASLD and MASH. QDs with a diameter of about 10 nanometers, comparable to the size of endogenous targets of SRs, are capable of emitting strong near-infrared (NIR) fluorescence. This NIR wavelength offers high tissue permeability and can be used for in vivo imaging analysis^21^. Furthermore, QDs are made water-soluble through chemical surface modification, mimicking metabolic waste products in the blood^28^. These properties make QDs well suited as a probe reporting SR-mediated clearance in the liver.

In this study, we report a non-invasive method to track the scavenger function of the hRES chronologically by use of QDs. We show that hepatic QDs retention declines with the progression from MASLD to MASH. This retention is mediated by SR-A on LSECs, and the decline is reversed upon disease regression. This approach provides a functional readout of liver pathology that is inaccessible to existing non-invasive tests, offering a possibility to overcome current challenges in MASH diagnosis.

## Results

### Quantum dots administered to the blood flow are sequestered in the liver

To investigate whether the physical properties of QDs could be utilized to non-invasively observe the liver in vivo, QDs were injected into the mice. Following intravenous administration of NIR-emitting QDs into mice, their distribution was monitored over time using an in vivo imaging system. The QDs began sequestration in the liver immediately after injection, and most of the QDs remaining in the body were captured within the liver after 72 hours (Fig. 1a). At the time point, the tissues were extracted and subjected to ex vivo imaging, showing that the fluorescence of QDs captured by the liver was more than 30-fold potent compared with other tissues (Fig. 1b, c).

**Fig. 1:**
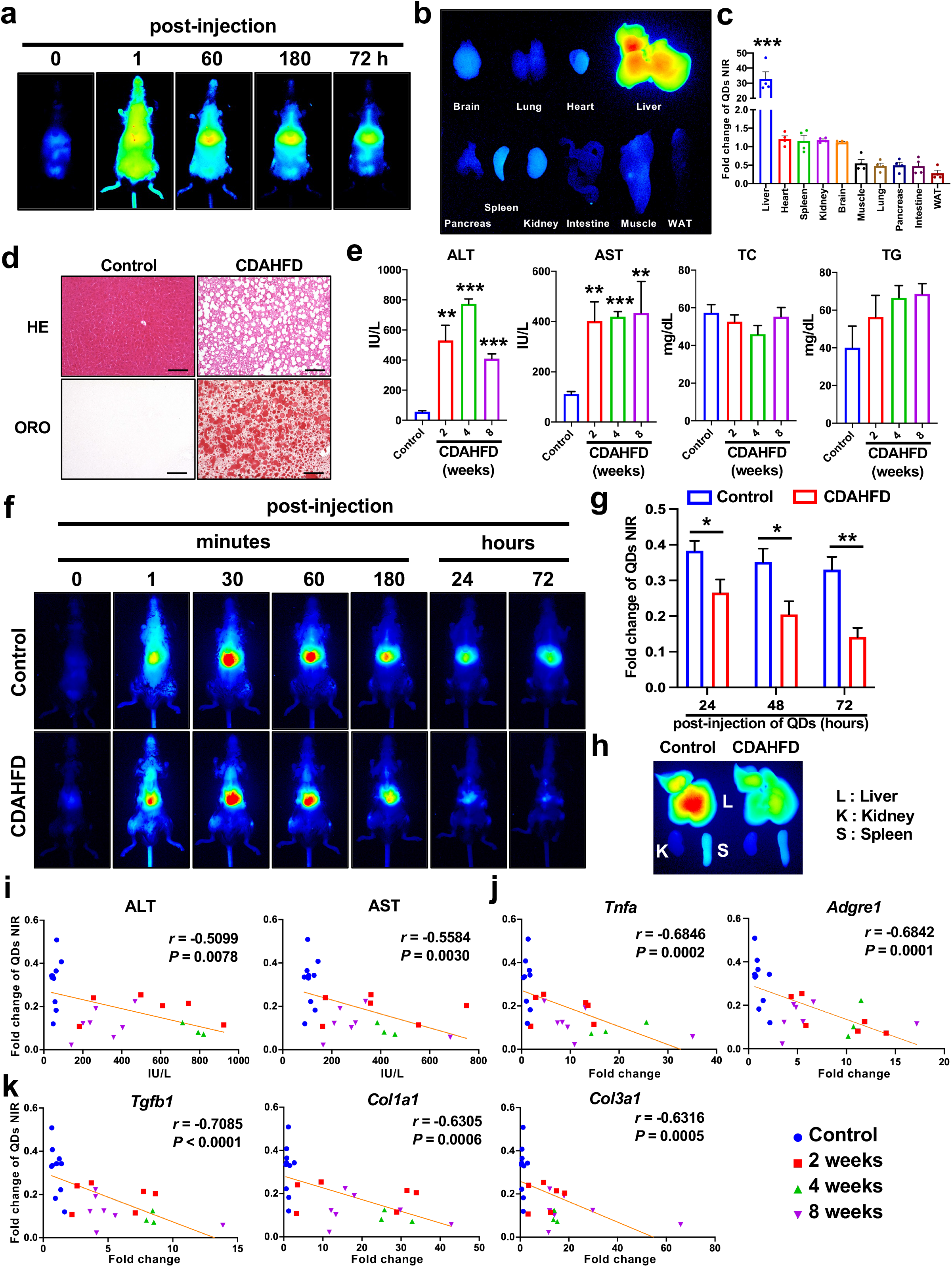
*in vivo* dynamics of QDs in normal and lean MASH model mice. **a**, Time-lapse in vivo images of QDs in a mouse after tail vein injection. **b**, Ex vivo observation of the tissues extracted from the mice 72 hours after QDs administration. **c**, Fluorescence intensity of QDs NIR in each extracted tissue. *n* = 4 mice; error bars, mean± s.e.m. \*\*\**P*<0.001, One-way ANOVA followed by Tukey’s test. **d**, Hematoxylin and eosin (HE) and oil red o (ORO) staining for frozen sections of control and CDAHFD-fed mouse liver. Scale bars, 100 μm. **e**, Serum concentration of alanine aminotransferase (ALT), aspartate aminotransferase (AST), total cholesterol (TC), and triacylglycerol (TG) in control and CDAHFD-fed mice. *n* = 8 (Control), *n* = 7 (2 weeks), *n* = 3 (4 and 8 weeks); error bars, mean± s.e.m. \*\**P*<0.01, \*\*\**P*<0.001. One-way ANOVA followed by Dunnett’s test. **f**, Time-lapse in vivo images of QDs in control and CDAHFD-fed mice. **g**, Fold change of QDs NIR of (f). *n* = 11 (Control), *n* = 7 (CDAHFD); error bars, mean± s.e.m. \**P*<0.05, \*\**P*<0.01. Two-way ANOVA followed by Tukey’s test. **h**, Ex vivo image of liver, kidney, and spleen extracted from the mice of (f). **i - k**, Scatter plots with linear trendline depicting the general relations between fold change of QDs NIR and serum hepatic injury markers (ALT and AST) (i), inflammatory marker genes (*Tnfa* and *Adgre1*) (j), and pro-fibrosis marker genes (*Tgfb1*, *Col1a1*, and *Col3a1*) (k). *n* = 9 (Control), *n* = 6 (2 weeks), *n* = 3 (4 weeks), *n* = 7 (8 weeks); *r* and *P* values were determined by spearman’s rank correlation coefficient.

### QDs sequestration in the liver is attenuated in the lean MASH model mice

Given that the blood clearance function of the liver is attenuated during MASH progression^5^, the QDs accumulation in the liver would be decreased. To clarify the hypothesis, MASH model mice followed by QDs administration were subjected to the in vivo imaging analysis. A choline-deficient, amino acid-defined high-fat diet (CDAHFD)-feeding is a well-established non-obese mouse MASH model^29, 30^. After eight weeks of CDAHFD-feeding, hematoxylin and eosin (HE) staining and Oil red O (ORO) staining for the liver sections showed marked lipid accumulation in the liver (Fig. 1d). The serum concentration of hepatic injury markers, alanine aminotransferase (ALT) and aspartate aminotransferase (AST) were markedly increased in the model independently of diet-feeding term, although serum total cholesterol (TC) and triglyceride (TG) were not significantly increased during the MASH progression (Fig. 1e). The tendency was consistent with previous reports^31^. After eight weeks of CDAHFD-feeding, mice were subjected to the QDs administration followed by in vivo imaging. In both the control and CDAHFD groups, hepatic accumulation of QDs peaked at 30 minutes post-injection and subsequently decreased over time (Extended Data Fig. 1a, b). When the fluorescence intensity at 30 minutes (the peak accumulation time) was normalized to 1.0, the relative QD fluorescence intensity in the CDAHFD group was significantly reduced compared to that in the control group from 24 hours post-injection (Fig. 1f, g). Ex vivo imaging analysis of the liver 72 hours after QDs administration revealed a marked decrease in QDs accumulation in MASH livers induced by CDAHFD, compared to that in the kidneys and spleen (Fig. 1h). To clarify the relationships between QDs accumulation and MASH pathogenesis, correlation analyses for serum hepatic injury markers (ALT and AST), inflammation (*Tnfa* and *Adgre1*), and fibrosis (*Tgfb1*, *Col1a1*, and *Col3a1*) gene expressions were performed. As previously reported^32, 33^, the severity of MASH had variability among individual mice, unrelated to the CDAHFD-feeding term. Nevertheless, the ALT and AST concentrations (Fig. 1i), hepatic *Tnfa* and *Adgre1* expressions (Fig. 1j), and hepatic *Tgfb1*, *Col1a1*, and *Col3a1* expressions (Fig. 1k) showed a potent negative correlation with QDs fluorescence.

### QDs accumulation in the liver is attenuated in the obesogenic MASH model mice

CDAHFD-feeding as the MASH model does not accompany obesity. In the clinical data, the ratio of lean MASLD patients is relatively higher in Asian countries, whereas the ratio is lower in Western countries^34^. The non-invasive evaluation for MASH was performed in the obesogenic model mice. A Gubra-Amylin NASH diet (GAND)-feeding, enriched with lipids and fructose, is one of the most prevalent obesogenic dietary MASH models. After 30 weeks of GAND-feeding, HE and ORO staining showed excess lipid accumulation in the liver (Fig. 2a). ALT and AST concentrations in the serum are significantly increased at 30 weeks of GAND-feeding, whereas they are not increased at four weeks of GAND-feeding. TC is significantly increased at four weeks of feeding, suggesting that four weeks of induction causes lipid metabolism dysfunction, resulting in MASLD but not MASH, whereas 30 weeks of induction cause inflammation like MASH (Fig. 2b). To observe the MASH pathology, QDs were injected into the GAND-fed mice. The NIR of the QDs was significantly attenuated at 72 hours after the injection of QDs in 30 weeks of GAND-induction. Whereas the accumulation was not significantly altered in four weeks of GAND-induction (Fig. 2c, d). The tissues were extracted and subjected to ex vivo imaging, demonstrating that four weeks of induction did not reduce QD accumulation in the liver, whereas 30 weeks of induction markedly reduced it (Fig. 2e). QDs NIR and hepatic injury markers, serum ALT and AST, showed a significant negative correlation (Fig. 2f). Pro-inflammatory genes (*Tnfa* and *Adgre1*) and pro-fibrogenic genes (*Tgfb1*, *Col1a1*, and *Col3a1*) expression also showed significant negative correlations with QDs NIR (Fig. 2g, h). These findings demonstrate that QDs-based monitoring is also applicable to observe the pathology of obesogenic MASH.

**Fig. 2:**
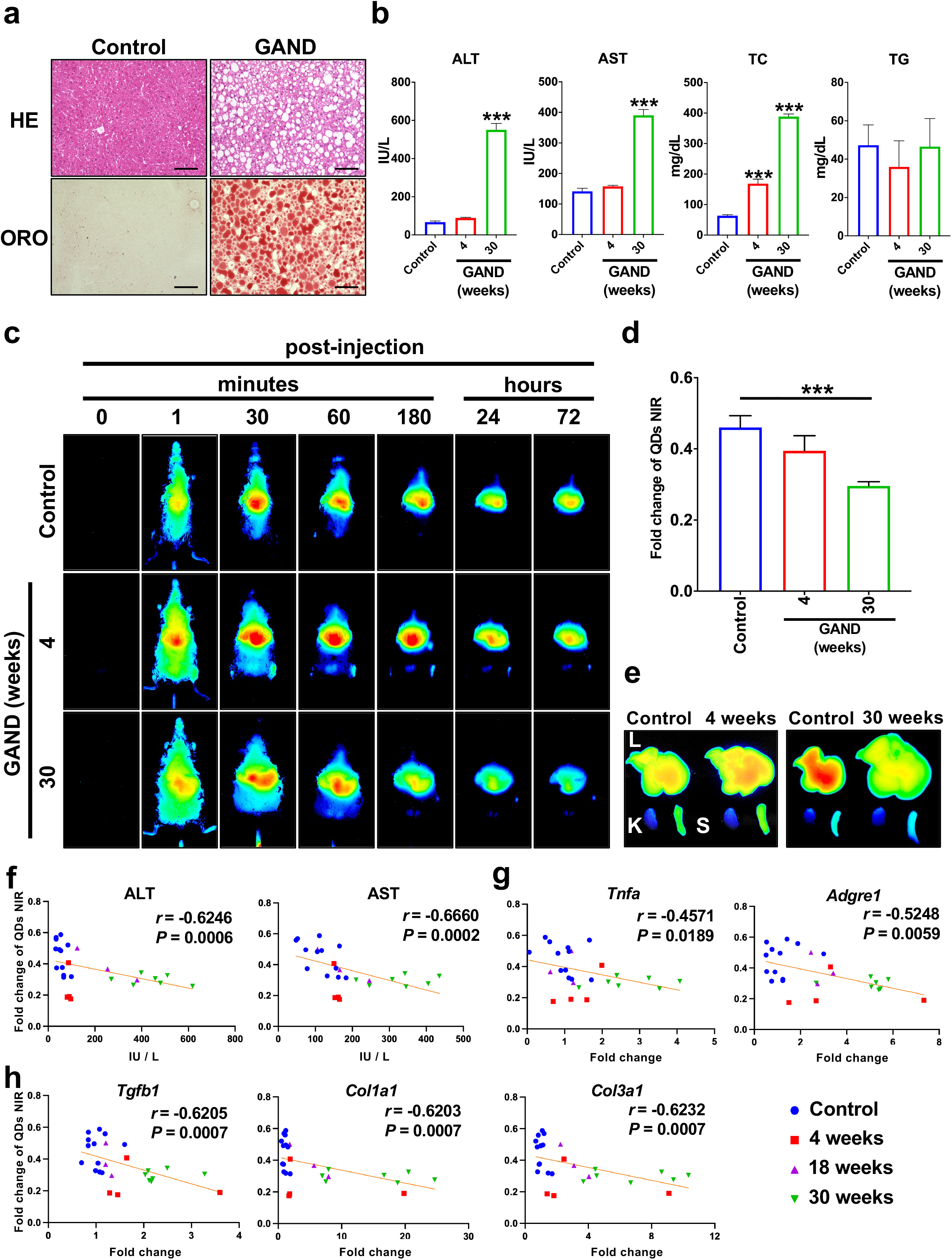
Hepatic QDs sequestration was attenuated in obesogenic MASH model mice. **a**, HE and ORO staining for frozen sections of control and GAND-fed mouse liver. Scale bars, 100 μm. **b**, Serum concentration of ALT, AST, TC, and TG in control and GAND-fed mice. *n* = 8 (Control), *n* = 4 (4 and 30 weeks); error bars, mean± s.e.m. \*\*\**P*<0.001. One-way ANOVA followed by Dunnett’s test. **c**, Time-lapse in vivo images of QDs in control and GAND-fed mice. **d**, Fold change of QDs NIR of (c). *n* = 10 (Control), *n* = 4 (4 weeks), *n* = 7 (30 weeks); error bars, mean± s.e.m. \*\*\**P*<0.001. One-way ANOVA followed by Tukey’s test. **e**, Ex vivo images of the liver, kidney, and spleen extracted from the mice of (d). **f - h**, Scatter plots with linear trendline depicting the general relations between fold change of QDs NIR and serum hepatic injury markers (ALT and AST) (f), inflammatory marker genes (*Tnfa* and *Adgre1*) (g), and pro-fibrosis marker genes (*Tgfb1*, *Col1a1*, and *Col3a1*) (h). *n* = 12 (Control), *n* = 4 (4 weeks), *n* = 3 (18 weeks), *n* = 7 (30 weeks); *r* and *P* values were determined by spearman’s rank correlation coefficient.

### Sequestered QDs reduction is observed in the injured liver but not in hepatic steatosis

To clarify which physiological functions in MASH pathology are involved in the decrease of QDs sequestration, analyses of an acute liver injury model induced by a single administration of Carbon Tetrachloride (CCl_4_) and a MASLD model by *ob/ob* mice were performed. A single administration of CCl_4_ induces severe pericentral oxidative stress, triggering necrosis in various cell types surrounding the central vein and temporal inflammation^35^. In contrast, the *ob/ob* mouse exhibits a MASLD-like phenotype characterized by marked hepatic lipid accumulation and impaired liver function. Conversely, because inflammation is not actively induced in the *ob/ob* model^36^, it serves as a valuable tool to analyze the relationship between QDs and lipid accumulation or hepatocyte injury independent of inflammatory processes. Time-lapse monitoring of serum ALT levels after a single CCl_4_ administration revealed a peak at 24 hours post-injection, at which point QDs were administered (Fig. 3a). HE staining showed extensive tissue necrosis in the peri-central area 24 hours after a single CCl_4_ injection. Whereas the liver exhibited marked regeneration at 96 hours, with only residual immune cells remaining around the central veins. No lipid deposition was detected at any time point during the process (Fig. 3b). Histological analysis of *ob/ob* mouse livers revealed marked lipid accumulation, while no tissue necrosis was observed (Fig. 3c). Serum biomarkers showed a significant increase in hepatocyte damage markers, ALT and AST, in addition to serum TC levels of *ob/ob* mice. The serum TG levels were not altered, consistent with a previous study^37^. At 96 hours after a single CCl_4_ administration, serum ALT and AST levels remained slightly but not significantly elevated compared with the control. Serum TC concentration was not altered, whereas serum TG was significantly increased (Fig. 3d). This tendency was also consistent with previous study^38^. Pro-inflammatory cytokine expressions (*Tnfa* and *Il1b*) in the livers showed a significant inhibition in *ob/ob* mice, whereas the expressions in the CCl_4_-treated mouse livers normalized 96 hours after the administration (Fig. 3e). Time-lapse monitoring of QDs NIR in the *ob/ob* and CCl_4_-administered mice uncovered that QDs sequestration is increased in *ob/ob* mice, whereas decreased in CCl_4_-administered mice (Fig. 3f, g). Ex vivo imaging of the livers also showed the same tendency (Fig. 3h). These data indicated that liver injury, rather than steatosis, significantly reduces hepatic QDs sequestration.

**Fig. 3:**
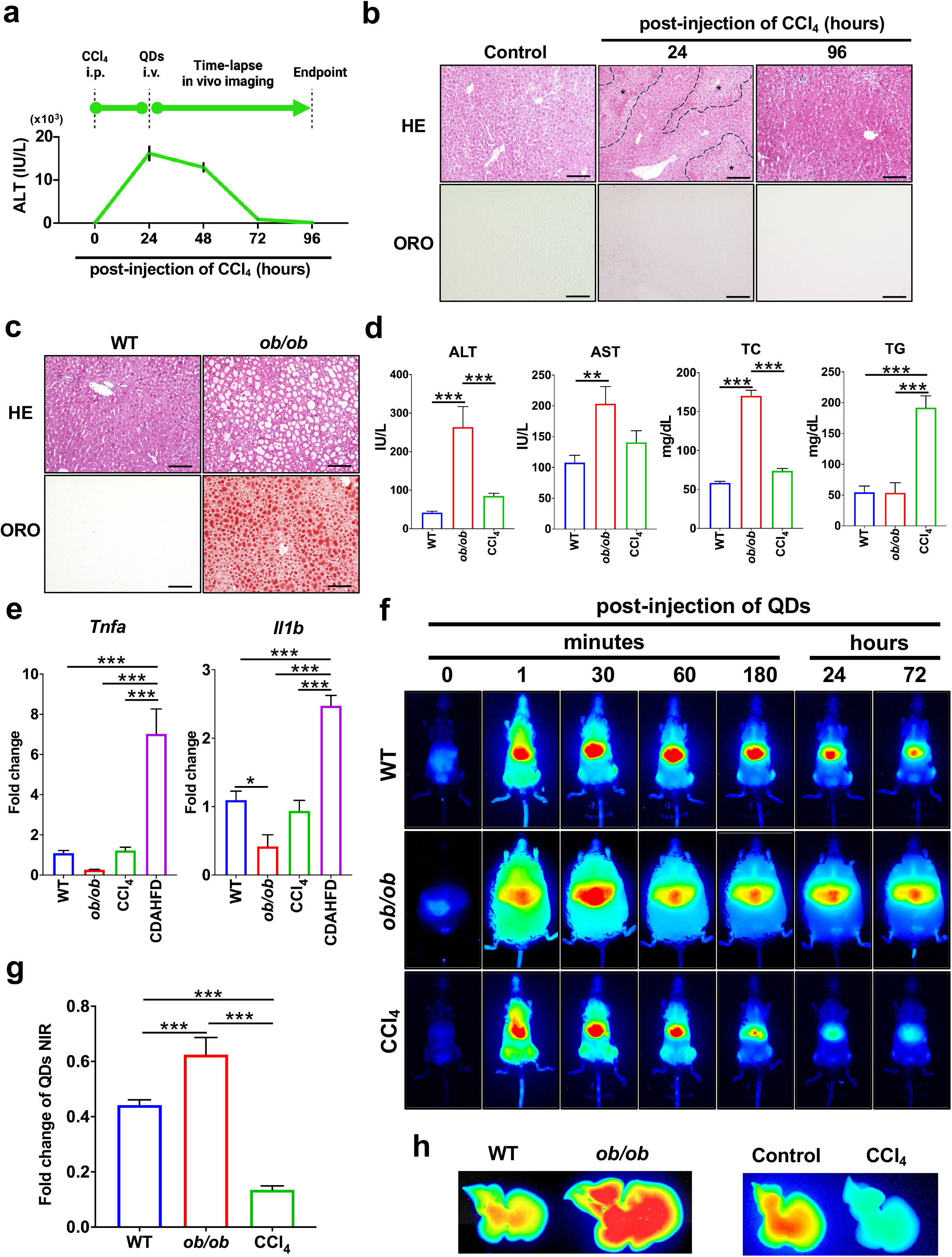
QDs sequestration is attenuated by CCl_4_-induced acute liver injury but not by steatosis or hepatocyte damage in *ob/ob* mice. **a**, Time course of CCl_4_ and QDs administration and ALT levels at the time points. **b**, HE and ORO staining of liver sections 24 and 96 hours after a single administration of CCl_4_ and the control. Asterisks surrounded by broken lines represent necrotic peri-central area. Scale bars, 100 μm. **c**, HE and ORO staining of liver sections from WT and *ob/ob* mice. **d**, Serum concentrations of ALT, AST, TG, and TC in WT, *ob/ob*, and CCl_4_-treated mice 96 hours post-administration. *n* = 15 (WT), *n* = 7 (*ob/ob*), n = 8 (CCl_4_); error bars, mean± s.e.m. \*\**P*<0.01. \*\*\**P*<0.001. One-way ANOVA followed by Tukey’s test. **e**, Fold change of pro-inflammatory cytokine expressions (*Tnfa* and *Il1b*) in WT (*n* = 12), *ob/ob* (*n* = 4), CCl_4_ (post-administration 96 h) (*n* = 8), CDAHFD (8 weeks) (*n* = 3); error bars, mean± s.e.m. \**P*<0.05. \*\*\**P*<0.001. One-way ANOVA followed by Tukey’s test. **f**, Time-lapse in vivo images of QDs in WT, *ob/ob*, and CCl_4_-administered mice. **g**, Fold change of QDs NIR of (f). *n* = 15 (WT), *n* = 7 (*ob/ob*), *n* = 8 (CCl_4_); error bars, mean± s.e.m. \*\*\**P*<0.001. One-way ANOVA followed by Tukey’s test. **h**, Ex vivo images of the liver extracted from the mice of (f).

### QDs are captured by LSECs affected by MASH progression

Given that necrotic injury, distinct from simple hepatocyte damage, is related to the reduction of QDs sequestration, the hepatic sinusoid could be directly involved in the sequestration mechanism. This hypothesis is consistent with our previous observation of the time course of sinusoidal injury and regeneration after a single CCl_4_ treatment^35^. Immunohistochemical staining for LSECs markers (Stab2 and FcγRⅡ/Ⅲ) in the liver sections showed sinusoidal destruction around the peri-central area at 24 hours after a single CCl_4_ injection (Extended Data Fig. 2a). qPCR analyses in the liver demonstrated that the fold change of QDs NIR has a significant negative correlation with pro-fibrogenic genes related to sinusoidal capillarization (*Tgfb1* and *Col1a1*) (Extended Data Fig. 2b). To identify the specific liver cells responsible for capturing administered QDs, mice were injected with QDs followed by FITC-labeled dextran. The livers were subsequently harvested, and the tissue sections were analyzed via microscopy. The analysis showed that QDs were captured within the hepatic sinusoid (Fig. 4a), suggesting that the cells consisting of hRES were responsible for the QDs capture. Immunohistochemical staining for the hepatic sections was performed to clarify the cells consisting of hRES. The analyses uncovered that both the CD31-positive LSECs and the F4/80-positive KCs were colocalized with QDs in the sinusoid (Fig. 4b). Flow cytometry (FCM) analysis for LSECs and KCs isolated by collagenase perfusion demonstrated that these cells took in the QDs (Fig. 4c). To determine whether LSECs or KCs are responsible for the decreased QDs uptake in MASH, QDs were administered to mice fed CDAHFD. Liver cells were isolated 30 minutes and 72 hours post-administration and subsequently analyzed by FCM. The analysis revealed that the significant reduction in QDs uptake under CDAHFD conditions occurred in LSECs, rather than in KCs (Fig. 4d). These results suggest that the impairment of scavenger function in LSECs, triggered by chronic inflammation, is responsible for the QDs uptake. Previous studies reported that the sinusoidal capillarization marker, CD34, was upregulated, whereas SR expression, including Stab2, was decreased in LSECs under the MASH condition^39–41^. Sinusoidal injury in the CDAHFD-induced MASH liver was evaluated by western blot analysis, elucidating that CD34 protein was significantly upregulated, whereas Stab2 was significantly decreased (Fig. 4e). Immunohistochemical staining for the markers also demonstrated CD34 upregulation and Stab2 downregulation in CDAHFD-induced MASH liver (Fig. 4f). Soluble ICAM-1 (sICAM-1) concentration in the serum is an endothelial injury marker, increased under MASH conditions^42, 43^. To analyze the LSECs damage, serum sICAM-1 concentration was measured, showing that both CDAHFD and GAND-inducible MASH resulted in a significant increase in sICAM-1, respectively (Fig. 4g, h), whereas *ob/ob* MASLD model mice did not show significant alteration (Fig. 4i). Healthy LSECs with robust scavenger functions actively facilitate material exchange through their fenestrated structure. Whereas, the LSECs undergo capillarization, accompanied by reduced expression of scavenger receptors and the loss of fenestrae under MASH conditions^40^. Liver sections of GAND-fed MASH mice were subjected to scanning electron microscopy (SEM) observation, showing that the abundant fenestration characteristic of healthy sinusoids in the control liver was largely diminished in the GAND-fed MASH mice (Fig. 4j and Extended Data Fig. 2c). These results demonstrated that inflammatory stimuli for LSECs impair scavenger function, reducing QDs sequestration in the liver.

**Fig. 4:**
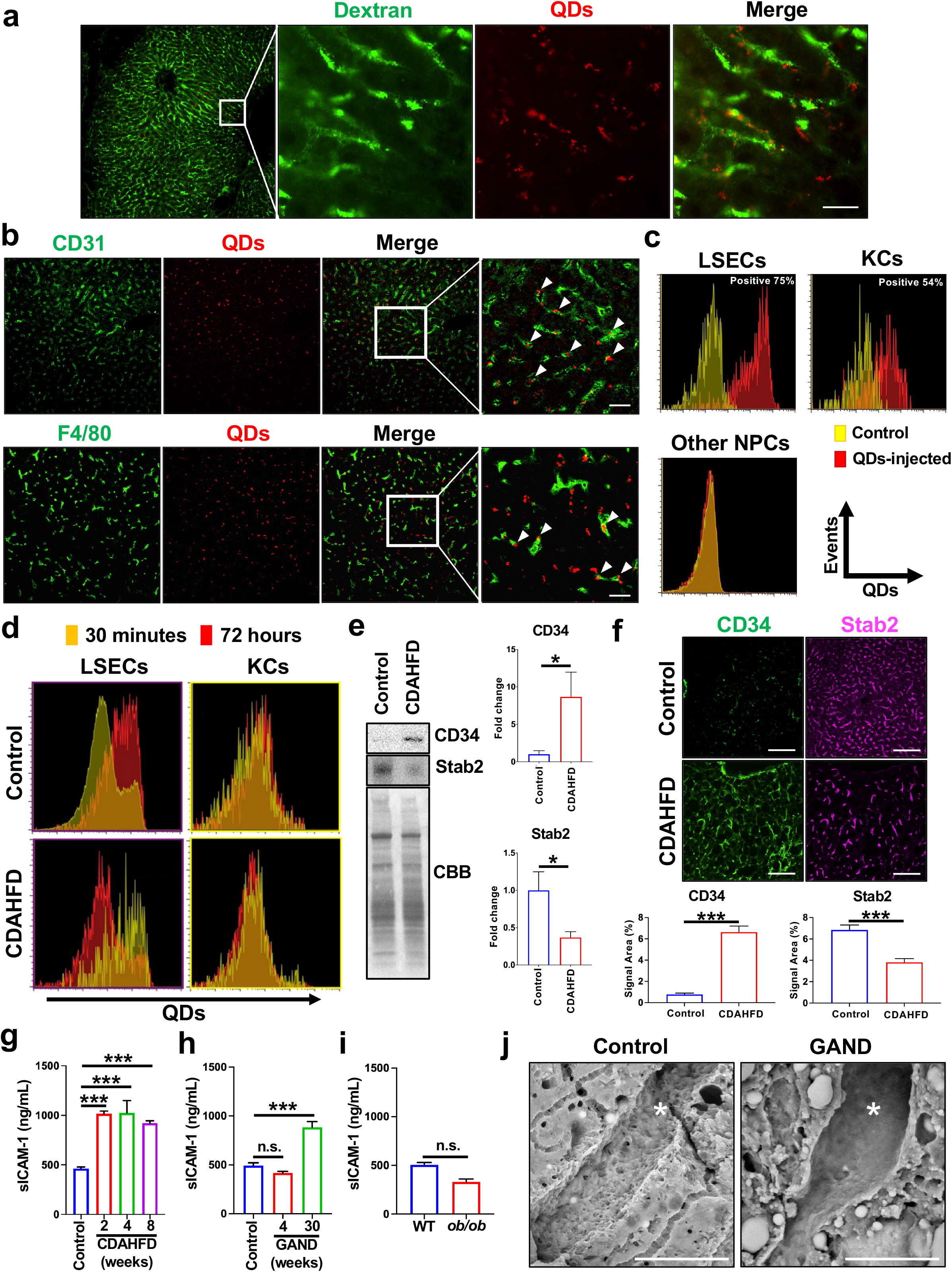
LSECs are the primary cells responsible for capturing QDs during MASH progression. **a**, Fluorescence imaging of hepatic sinusoids in QDs and FITC-labeled dextran-administered mouse liver section. Scale bar, 20 μm. **b**, Immunohistochemical staining of CD31 and F4/80 for QDs-administered mouse liver sections. Scale bars, 20 μm. **c - d**, FCM analysis of primary hepatic non-parenchymal cells (NPCs) isolated from control and QDs-administered mouse livers via collagenase perfusion. Characterization of LSECs and KCs 24 hours after QDs injection (c). Comparative analysis of QDs uptake by hepatic LSECs and KCs isolated from the livers of control and CDAHFD-fed mice at 30 minutes and 72 hours after QDs administration (d). **e**, Western blot analysis for CD34 and Stab2 in the control and CDAHFD-fed mouse livers. *n* = 7 (Control and CDAHFD); error bars, mean± s.e.m. \**P*<0.05. Welch’s t-test. **f**, Immunohistochemical staining of CD34 and Stab2 in control and CDAHFD-fed mouse livers. Scale bars, 100 μm. *n* = 8 (Control and CDAHFD); error bars, mean± s.e.m. \*\*\**P*<0.001. Welch’s t-test. **g - i**, Serum concentration of soluble ICAM-1 (sICAM-1) in MASLD/MASH model mice. Control (*n* = 11), CDAHFD 2 weeks (*n* = 7), 4 weeks (*n* = 3), and 8 weeks (*n* = 7) (g). Control (*n* = 11), GAND 4 weeks (*n* = 4), and 30 weeks (*n* = 7) (h). WT (*n* = 3) and ob/ob (*n* = 3) (i). Error bars, mean± s.e.m. \*\*\**P*<0.001. One-way ANOVA followed by Tukey’s test. **j**, Scanning electron microscopy observation of Control and GAND-fed mouse liver sections. Asterisks represent hepatic sinusoids. Scale bars, 10 μm.

### QDs sequestration by LSECs is mediated by scavenger receptor class A

To explore the mechanism of QDs sequestration by LSECs, competitive inhibitors for SRs were injected before QDs administration. A previous study reported that the scavenger receptor Stab2 mediates the QDs sequestration in Zebrafish^44^, suggesting that Stab2 inhibition could prevent QDs uptake in LSECs. Dextran sulfate (DS) is a negatively charged polyanionic polysaccharide, potently binding to multiple SRs. Hyaluronic acid (HA) and heparin are selective ligands for Stab2, resulting in competitive inhibition of Stab2. DS, HA, and heparin were intravenously injected into the mice, followed by QDs administration. The in vivo imaging analyses intriguingly showed that DS, but not HA or heparin, remarkably prevented QDs sequestration, suggesting that SRs other than Stab2 mediated the QDs uptake by LSECs (Fig. 5a, b, c, and Extended Data Fig. 3a). QDs reduction by DS was observed not only in the liver but also in the spleen (Fig. 5d), indicating that the scavenger receptor related to QDs sequestration is potently but not exclusively expressed in LSECs. Scavenger receptor class A (SR-A) is a scavenger receptor family that has five family members known as SR-A1, -A3, -A4, -A5, and -A6^45^. SR-A1, having a high affinity to DS, is abundantly expressed in LSECs, and plays a significant role in capturing harmful lipids, such as oxidized LDL and acetylated LDL^40^. These LDL accumulations in the blood significantly elevate hepatic lipotoxicity, resulting in MASH aggravation and a higher risk of hepatic sclerosis and cardiovascular events^46, 47^. To determine whether SR-A is responsible for the QDs sequestration, QDs were administered into SR-A1 and SR-A6 double-knockout (SR-A1/6 KO) mice, followed by time-lapse in vivo imaging. The analysis demonstrated that QDs sequestration was potently attenuated in the SR-A1/6 KO mouse livers (Fig. 5a, b, c, d). LSECs were isolated from the WT or SR-A1/6 KO mouse livers by collagenase perfusion followed by primary culture and FCM analyses (Fig. 5e). Tannic acid (TA), a competitive inhibitor of SR-A, was administered to primary cultured LSECs in addition to DS. These primary cultured LSECs were treated with QDs, then subjected to immunocytochemistry and FCM analyses. Immunocytochemistry showed that the amount of QDs taken by CD31 and Stab2 double-positive LSECs was decreased by DS or TA administration. The reduction was also observed in SR-A1/6 KO LSECs (Fig. 5f). FCM analyses revealed that DS (Fig. 5g) and TA (Fig. 5h) treatments, and SR-A1/6 KO (Fig. 5i) markedly reduced QDs sequestration in the LSECs. These results demonstrated that SR-A expressed in LSECs is the responsible scavenger receptor for QDs sequestration.

**Fig. 5:**
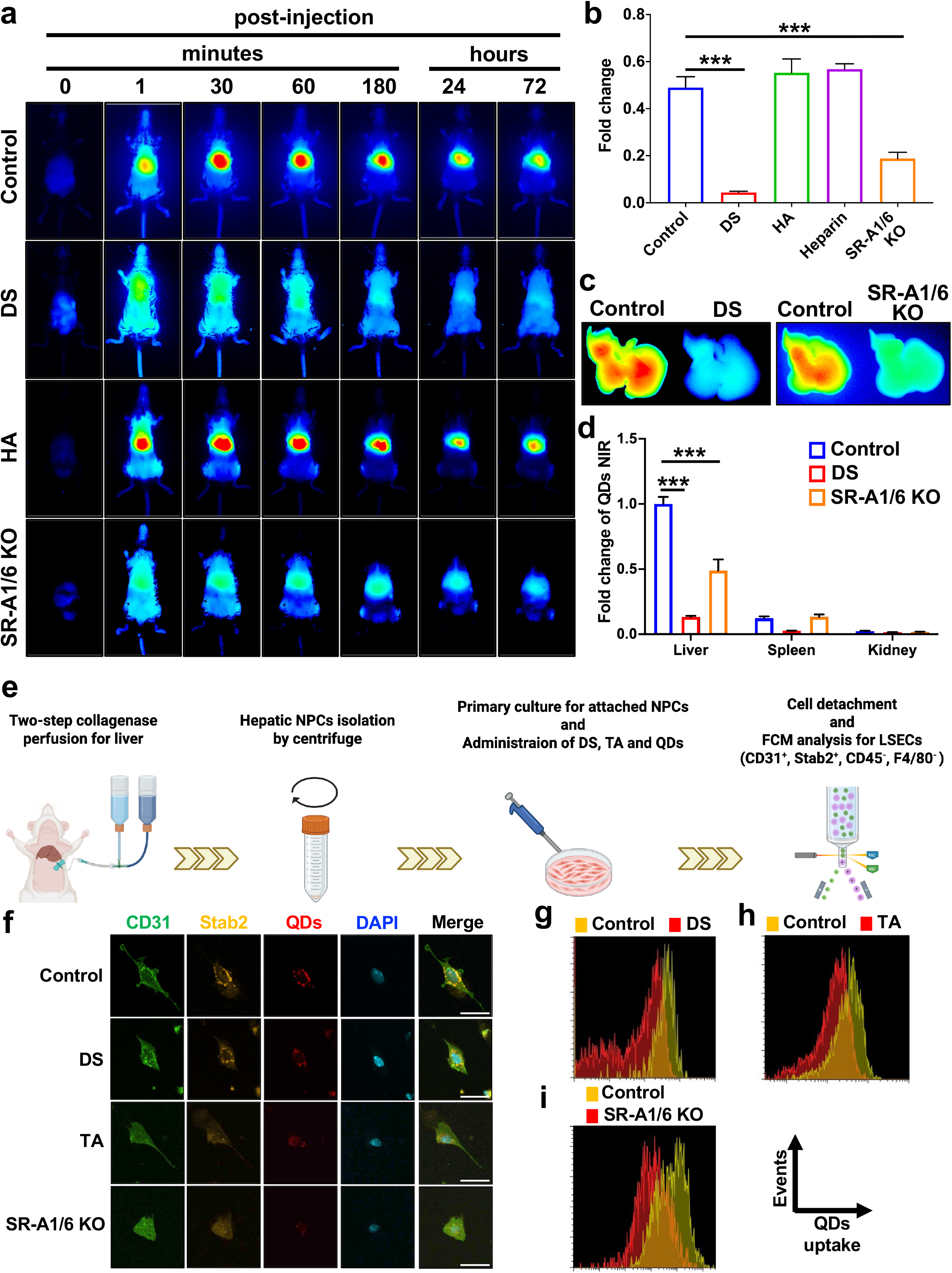
SR-A expressed in LSECs is responsible for QDs sequestration. **a**, Time-lapse in vivo images of QDs in control, dextran sulfate (DS), hyaluronic acid (HA) -treated, or SR-A1/6 knockout mice after QDs injection. **b**, Fold change of QDs NIR of (a). *n* = 7 (Control), *n* = 5 (DS), *n* = 3 (HA), *n* = 5 (SR-A1/6 KO); error bars, mean± s.e.m. \*\*\**P*<0.001. One-way ANOVA followed by Dunnett’s test. **c**, Ex vivo image of livers extracted from control, DS-treated, and SR-A1/6 KO mice. **d**, Fluorescence intensity of QDs NIR of liver, spleen, and kidney extracted from each mouse. *n* = 4 (Control), *n* = 5 (DS), and *n* =5 (SR-A1/6 KO); error bars, mean± s.e.m. \*\*\**P*<0.001, One-way ANOVA followed by Dunnett’s test. **e**, Scheme of primary LSECs isolation and flow cytometry (FCM) analysis. **f**, Immunohistochemical staining of CD31 and Stab2 for primary cultured LSECs. Before QDs administration, the LSECs were treated with DS or TA, or isolated from SR-A1/6 KO mice. Scale bar, 20 μm. **g – i**, FCM analysis for LSECs treated with DS (g), TA (h), and isolated from SR-A1/6 KO mice (i).

### QDs sequestration in the LSECs sensitively reflects the pathological condition of MASH

Given that non-invasive evaluation of MASH reversibility is achieved using QDs, it serves as a valuable tool for establishing surrogate endpoints in MASH drug development. Furthermore, it clarifies the “point of no-return” in assessing disease progression. It is known that the expression of SRs in LSECs decreases with the progression of MASH and recovers during sinusoidal regeneration^40^. To investigate whether MASH progression and recovery could be evaluated non-invasively by QDs, a CDAHFD-induced MASH model with a recovery term was subjected to in vivo imaging analysis (Fig. 6a). Two weeks of recovery remarkably restored the hepatic injury markers ALT and AST, whereas the serum lipid concentrations did not change (Fig. 6b). QDs were administered to the control, CDAHFD, and recovery groups, and their accumulation in the liver was analyzed over time via in vivo imaging. The results demonstrated that QDs accumulation in the recovery group significantly improved starting at 24 hours post-injection compared to the CDAHFD group (Fig. 6c, d, e). Correlation analyses between serum liver injury markers and QDs NIR fluorescence showed that both ALT and AST were significantly and negatively correlated with the QDs NIR (Fig. 6f). In addition, a negative correlation was observed with the expression level of pro-inflammatory and pro-fibrogenic genes (Fig. 6g, h). These results demonstrated that under MASH conditions, it is possible to evaluate not only the exacerbation of hRES dysfunction but also its subsequent recovery (Fig. 6i and Extended Data Fig. 3).

**Fig. 6:**
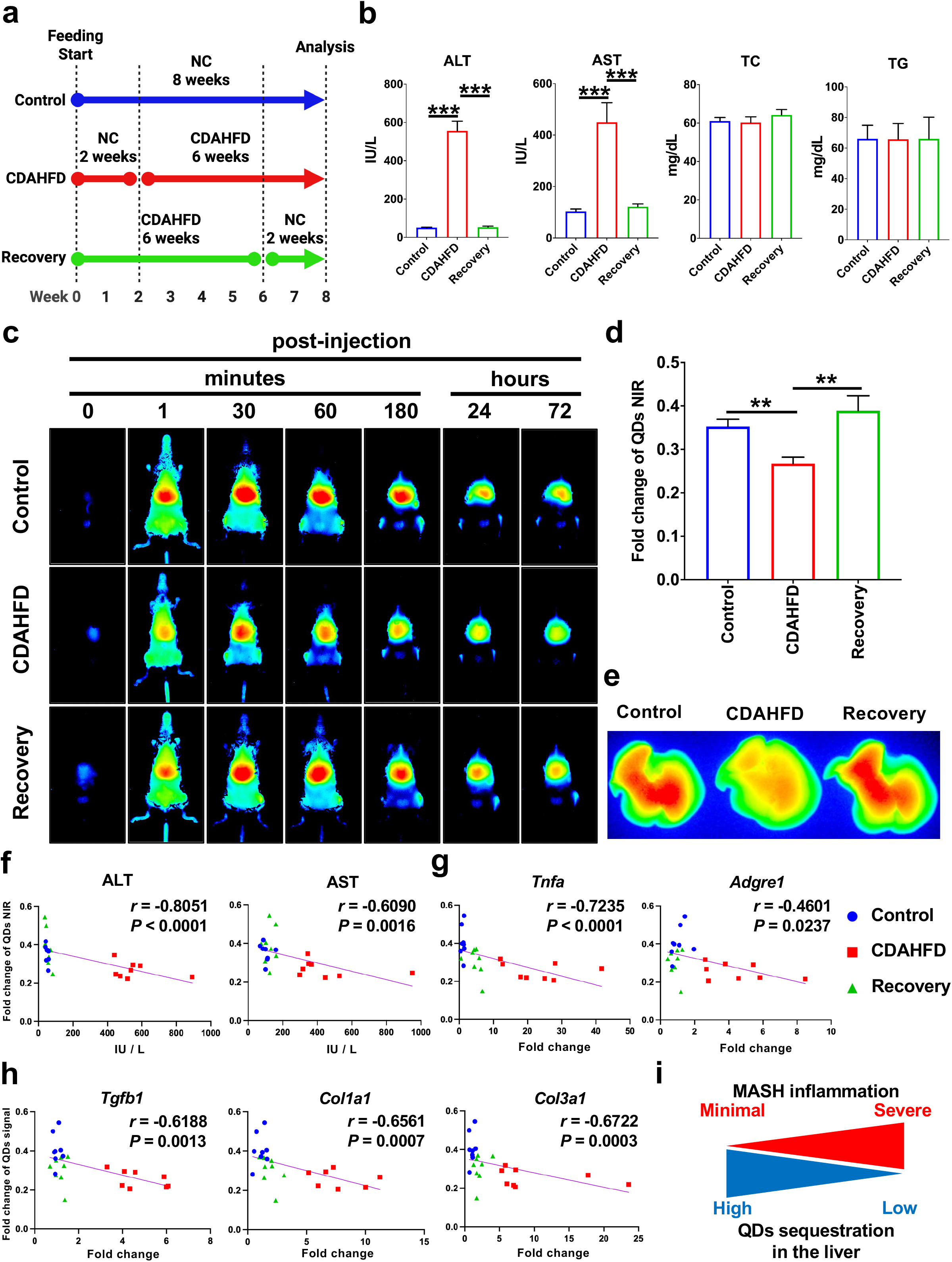
QDs NIR sensitively reflect hRES dysfunction and regeneration. **a,** Time-course of CDAHFD feeding in each group. **b**, Serum concentration of ALT, AST, TC, and TG in control, CDAHFD, and recovery group. *n* = 8 (Control, CDAHFD, and Recovery); error bars, mean± s.e.m. \*\*\**P*<0.001. One-way ANOVA followed by Tukey’s test. **c**, Time-lapse in vivo images of QDs in control, CDAHFD, and recovery group. **d**, Fold change of QDs NIR of (c). *n* = 8 (Control, CDAHFD, and Recovery); error bars, mean± s.e.m. \**P*<0.05, \*\**P*<0.01. Two-way ANOVA followed by Tukey’s test. **e**, Ex vivo images of the liver extracted from the mice of (d). **f - h**, Scatter plots with linear trendline depicting the general relations between fold change of QDs NIR and serum hepatic injury markers (ALT and AST) (f), inflammatory marker genes (*Tnfa* and *Adgre1*) (g), and pro-fibrosis marker genes (*Tgfb1*, *Col1a1*, and *Col3a1*) (h). *n* = 8 (Control, CDAHFD, and Recovery); *r* and *P* values were determined by spearman’s rank correlation coefficient. **i**, Scheme of inverse correlation between MASH inflammation and QDs sequestration that the current study demonstrated.

## Discussion

In this study, we demonstrate that the scavenger function of LSECs can be monitored non-invasively and longitudinally in MASH by leveraging the optical and material properties of QDs. Conventional approaches to assess this function, such as two-photon microscopy, require liver exteriorization and thus preclude real-time, longitudinal tracking during disease progression and recovery. In contrast, spatiotemporal imaging using QDs overcomes these constraints and captures the dynamic alterations of the scavenger function of LSECs in the same animal. Unlike currently available non-invasive tests, which report circulating biochemical parameters or gross structural alterations of the liver, this approach reads out cellular function within the tissue.

Unlike the immediate reduction in hepatic QDs entrapment caused by SR-A inhibition or knockout, QDs retention in MASH livers decayed gradually over time. This kinetic shift, specifically in peak timing, indicates that reduced QDs signal in MASH reflects impaired LSECs scavenger performance, including restricted fenestral transport, rather than a mere loss of LSECs or SR-A expression. Supporting this, SEM revealed fenestral loss in MASH sinusoids, which impairs particle access to LSECs independently of receptor abundance. Thus, QDs kinetics report the functional state of LSECs beyond single-molecule expression.

To define what the QDs signal reports, we compared models that dissociate steatosis from injury. Hepatic QDs retention was markedly reduced after a single CCl_4_ administration, whereas it was preserved, and even increased, in *ob/ob* mice, which exhibit pronounced steatosis without active inflammation. The signal therefore reflects sinusoidal injury rather than lipid accumulation, which addresses the clinical need to distinguish MASLD from MASH. On the other hand, because the signal declined in CCl_4_-induced acute injury, and correlated comparably with inflammatory and pro-fibrogenic gene expression, this approach is not specific to MASH. Defining which LSEC alterations, such as capillarization, fenestrae loss, or reduced SR-A expression, underlie the reduced QDs uptake remains an important future task.

Translating this approach to humans requires addressing optical penetration and material biosafety. First, the 820-nm NIR fluorescence used here traverses the few millimeters between the body surface and the liver in mice, whereas the human liver lies several centimeters deep. The present configuration therefore cannot be applied directly to patients. What this study establishes, however, is a principle rather than a particular reporter: the functional state of LSECs can be readout from the receptor-mediated clearance of administered particles. This principle may be transferable to modalities with greater tissue penetration, including NIR-II/short-wave infrared imaging, photoacoustic imaging, and nuclear medicine probes targeting SR-A. Second, conventional cadmium-based QDs used here raise concerns about heavy metal toxicity and long-term retention in the body. cadmium-free alternatives, including silver indium sulfide, silver germanium sulfide, carbon quantum dots, and multinary quantum dots are under development^48^. Such next-generation cadmium-free QDs improve biosafety while allowing precise control of particle size and surface biocompatibility, thereby enabling tailored pharmacokinetics. NIR-II–emitting QDs minimize autofluorescence and enable deep-tissue imaging. Developing such materials is a prerequisite for clinical translation.

Several limitations should be noted. First, the QDs signal was not compared with histological scoring. Disease severity was assessed using serum transaminases and the hepatic gene expression. The relationship between QDs retention and established histological indices remains to be determined. Second, our findings were obtained in mice. The SR-A dependence requires validation in human LSECs, given species differences in scavenger receptor expression and sinusoidal architecture.

In conclusion, we established a non-invasive method to visualize the scavenger function of LSECs in living mice using QDs. The signal declined with MASH progression, depended on SR-A, and recovered upon disease regression. This approach provides a functional readout of the liver that is inaccessible to existing non-invasive assessments, offering a new basis for evaluating disease activity and therapeutic response in MASH.

## Methods

### Animal models

To establish the control group, eight-to-ten-week-old wild-type C57BL/6N male mice (Japan SLC, Shizuoka, Japan) were maintained on a standard chow diet (CE-2; CLEA Japan, Tokyo, Japan). To induce metabolic dysfunction-associated steatohepatitis (MASH), mice were fed either a choline-deficient, L-amino acid-defined, high-fat diet (CDAHFD; A06071302, Research Diets, New Brunswick, NJ, USA) *ad libitum* for 2, 4, or 8 weeks, or a high-fat, high-fructose, and high-cholesterol diet, also known as Gubra Amylin NASH diet, (GAND; D09100310, Research Diets) *ad libitum* for 4, 18, or 30 weeks. As a non-MASH model of metabolic dysfunction-associated steatotic liver disease (MASLD), eight-week-old male *ob/ob* mice (C57BL/6JHamSlc-*ob/ob*; Japan SLC) and their wild-type littermates (C57BL/6JHamSlc-+/+; Japan SLC) were utilized. SR-A1/6 KO mice^49, 50^ were kindly provided by K. Tryggvason (Duke-NUS Medical School).

### In vivo imaging and quantum dots administration

For in vivo imaging of control and MASH/MASLD model mice, carboxylated quantum dots (QDs; Q21371MP, Thermo Fisher Scientific, Waltham, MA, USA) diluted in phosphate-buffered saline (PBS) were administered via the tail vein at a dose of 2.4 pmol/g body weight. Following the tail vein injection of QDs, the *in vivo* distribution was monitored over time using the Pearl Trilogy imaging system (LI-COR Biosciences, Lincoln, NE, USA). Intermittent imaging was performed from pre-injection (0 min) up to 72 h post-injection, detecting near-infrared fluorescence at 820 nm excited by a 785 nm laser. To quantify the signal intensity of QDs in the liver, a region of interest (ROI) was defined over the area of maximum signal intensity in the liver, while the inguinal region—where no QD accumulation was observed—was used to measure the background signal. The absolute fluorescence intensity of the liver ROI was then determined. For temporal analysis, the relative signal intensity at each time point was calculated by normalizing the values to the peak signal intensity observed at 30 min post-injection, which was defined as one.

### Anesthesia, euthanasia, and tissue preparation

Following the completion of *in vivo* imaging, mice were anesthetized with isoflurane to collect blood samples, and then humanely euthanized by cervical dislocation. Organs were immediately harvested for subsequent *ex vivo* imaging, quantitative reverse transcription-polymerase chain reaction (qPCR), and histological and biochemical analyses. For multi-organ *ex vivo* analysis, mice were euthanized via carbon dioxide inhalation, immediately followed by transcardial perfusion with PBS and 4% paraformaldehyde (PFA). The harvested organs were then post-fixed by immersion in 4% PFA.

### Quantitative RT-PCR (qPCR)

Total RNA was extracted from the harvested mouse livers using Sepasol-RNA I Super G (09379-55; Nacalai Tesque, Kyoto, Japan) according to the manufacturer’s protocol. The isolated total RNA was then reverse-transcribed to synthesize cDNA using the ReverTra Ace qPCR RT Master Mix with gDNA Remover (FSQ-301; Toyobo, Osaka, Japan). qPCR was performed on a LightCycler 96 system (Roche Diagnostics, Basel, Switzerland) using the KAPA SYBR FAST qPCR Kit (KK4611; Nippon Genetics, Tokyo, Japan) and gene-specific primer sets. The relative expression levels of the target genes were calculated using the comparative threshold cycle (ddCt) method, with glyceraldehyde-3-phosphate dehydrogenase (*Gapdh*) utilized as the internal housekeeping gene for normalization. Primers used in this study are listed in Supplementary Table S1.

### Biochemical analysis of serum

Blood samples collected from the mice used for *in vivo* imaging were allowed to stand at room temperature for 30 minutes to facilitate clotting. The samples were then centrifuged at 1,000 × *g* for 20 minutes to separate the serum from the blood clot. The recovered serum was analyzed to measure alanine aminotransferase (ALT), aspartate aminotransferase (AST), triglyceride (TG), and total cholesterol (TC) levels using a Fuji Dry-Chem NX700 analyzer (Fujifilm, Tokyo, Japan) with the corresponding Fuji Dry-Chem Slides for GPT/ALT, GOT/AST, TG, and TCHO (Fujifilm, Tokyo, Japan).

### Histological and immunohistochemical analyses

Harvested liver tissues were sectioned into small pieces and embedded in Tissue-Tek O.C.T. Compound (4583; Sakura Finetek, Tokyo, Japan) using Cryomold molds (4566; Sakura Finetek). The embedded frozen blocks were stored at -80°C. For sectioning, the blocks were cut into 6-μm-thick sections using a cryostat maintained at -20°C. The cryosections were mounted on glass slides, air-dried, and fixed with 4% PFA. Hematoxylin (Fujifilm Wako Pure Chemical, Osaka, Japan) and eosin (Nacalai Tesque, Kyoto, Japan) (H&E) staining, as well as Oil Red O (Nacalai Tesque, Kyoto, Japan) staining, were performed according to standard protocols. For immunohistochemical staining, nonspecific binding was blocked with 5% skim milk in PBS. The sections were then incubated with primary antibodies, including an anti-CD31 antibody (ab28364; Abcam, Cambridge, UK), anti-F4/80 antibody (MCA497GA; Bio-Rad, CA, USA), anti-CD34 (54158; Cell Signaling Technology, MA, USA), anti-FcγRⅡ/Ⅲ (553141; BD Biosciences, NJ, USA) and anti-Stab2 (D317-3; Medical and Biological Laboratories, Tokyo, Japan). After the incubation, sections were rinsed and incubated with secondary antibodies, followed by DAPI staining.

### Isolation of hepatic non-parenchymal cells

Following the tail vein administration of QDs, normal or MASH model mice were euthanized via carbon dioxide inhalation. Immediately after euthanasia, the abdomen was opened, and hepatic cells were isolated using the collagenase perfusion method, in which collagenase Type X (039-17864; Fujifilm Wako Pure Chemical) was perfused through the portal vein. To separate the non-parenchymal cell (NPC) fraction, the resulting liver cell suspension was subjected to differential centrifugation: initially at 50 × *g* once, and subsequently at 70 × *g* three times. The pellet containing parenchymal hepatocytes was discarded after each centrifugation step, leaving the NPCs in the supernatant. The isolated NPC fraction was then incubated with red blood cell (RBC) lysis buffer to eliminate erythrocytes. To prevent non-specific antibody binding, Fc receptors were blocked using TruStain FcX (101319; BioLegend, San Diego, CA, USA) prior to antibody staining.

### Flow cytometry and cell sorting

For the detection and quantification of QD uptake, the isolated NPCs were stained with the following antibodies and reagents: PerCP/Cy5.5-conjugated anti-CD45 antibody (103131; BioLegend), PE/Cy7-conjugated anti-F4/80 antibody (123113; BioLegend), APC-conjugated anti-CD31 antibody (102409; BioLegend), Alexa Fluor 488-conjugated anti-Stab2 antibody (D317-A48; Medical and Biological Laboratories) and DAPI (19178-91; Nacalai Tesque) for viability staining. The fluorescence signals of QDs within LSECs and KCs were subsequently detected and analyzed using an SH800S cell sorter (Sony Biotechnology, San Jose, CA, USA).

### Tissue preparation for scanning electron microscopy

Mice were deeply anesthetized with isoflurane and transcardially perfused through the left ventricle using a peristaltic pump at a flow rate of approximately 1 mL/min, with the right atrium incised for drainage. After perfusion with PBS for 1–2 min to remove blood, the fixative was gradually added to the PBS perfusate without interrupting the flow. PBS and fixative were initially mixed at a 1:1 ratio, and the proportion of fixative was progressively increased to full strength (1.5% paraformaldehyde and 3% glutaraldehyde in 0.1 M phosphate buffer, pH 7.4). Perfusion fixation was completed within approximately 5–10 min.

The liver was excised, trimmed, and sectioned into 200-µm-thick slices using a vibrating microtome (LinearSlicer, Dosaka EM, Kyoto, Japan). The slices were immersed in the same fixative for about 30 min, rinsed with distilled water, and post-fixed with 1% osmium tetroxide (OsO₄) on ice for 30 min. After rinsing with distilled water, the specimens were dehydrated with ethanol and air-dried without critical point drying. The dried specimens were sputter-coated with platinum and examined using a Phenom Pro desktop scanning electron microscope (Thermo Fisher Scientific) at an accelerating voltage of 10 kV.

## Acknowledgements

We thank Frank J. Gonzalez for advice on the research plan and proofreading. We thank Mitsuko Kobayashi, Satomi Ohnishi, Yusuke Konishi, and Fumitaka Okubo for assistance with the experiments. We thank The Center for Animal Research and Education (CARE) at Nagoya University for technical supports for animal experiments. This project was funded by JSPS KAKENHI 25K14934 (TY), 24K21339 (HY), 25H01217 (HY), 24KK0150 (TS) and 26K02216 (TS), the Japan Agency for Medical Research and Development (LEAP, [26gm0010011s0103] and Research Program on Hepatitis, [26fk0210178h0002]) (TS), the MEXT Quantum Leap Flagship Program (MEXT Q-LEAP, JPMXS0120330644) (HY), and Grants-in-aid of NAKATANI Foundation (TY), The HORI Science and Arts Foundation (TY, TS), SEI Group CSR Foundation, SECOM Science and Technology Foundation (HY, TS), and Takeda Science Foundation (TS).

## Author contributions

T.Y., M.M., S.S., E.W., H.K., K.O., I.S., and X.Y. performed the research and analyzed the data. T.Y., A.I., M.T., S.A., Y.B., H.Y., and T.S. designed and supervised the research. T.Y., H.Y., and T.S. wrote the manuscript.

## Competing interests

The authors declare no competing financial interests.

---

**Extended Data Fig. 1:**
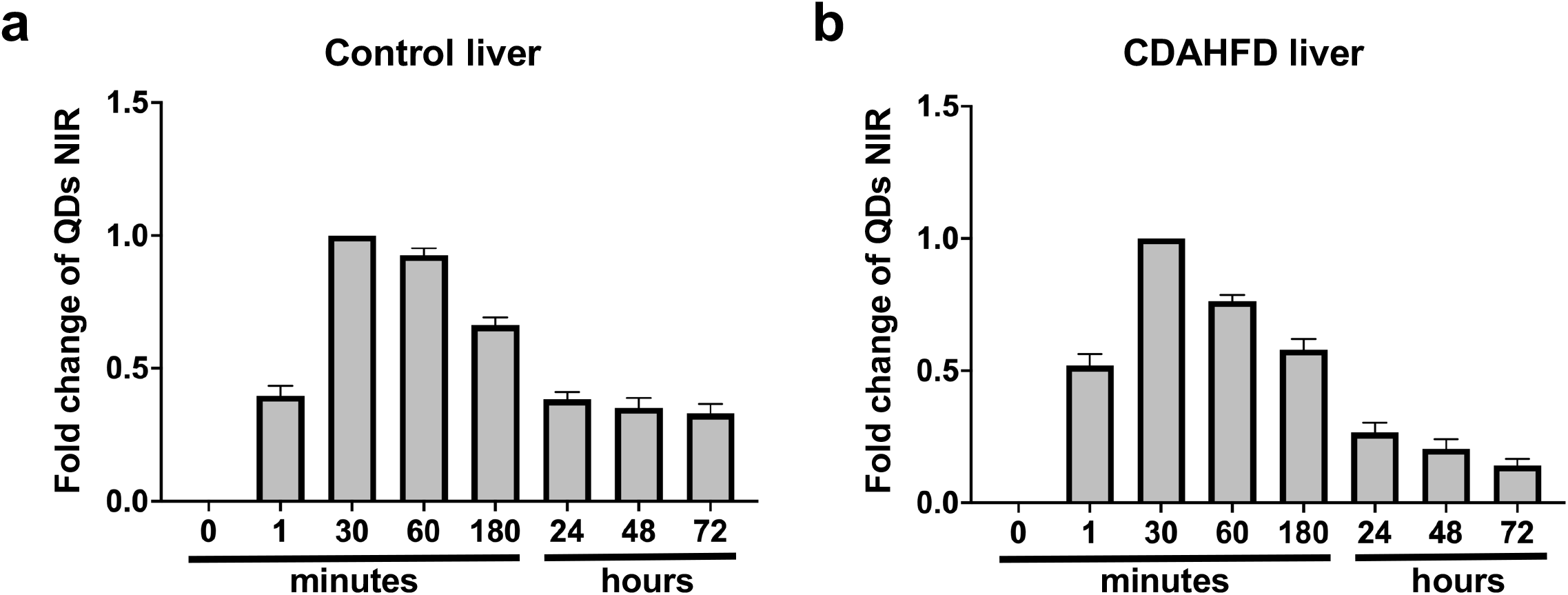
Time-lapse alteration of ǪDs NIR fluorescence detected by in vivo imaging system. **a - b**, The time-dependent relative fluorescence intensity in the normal (a) and CDAHFD-fed (b) mouse livers, normalized to the value obtained 30 minutes after ǪDs administration.

**Extended Data Fig. 2:**
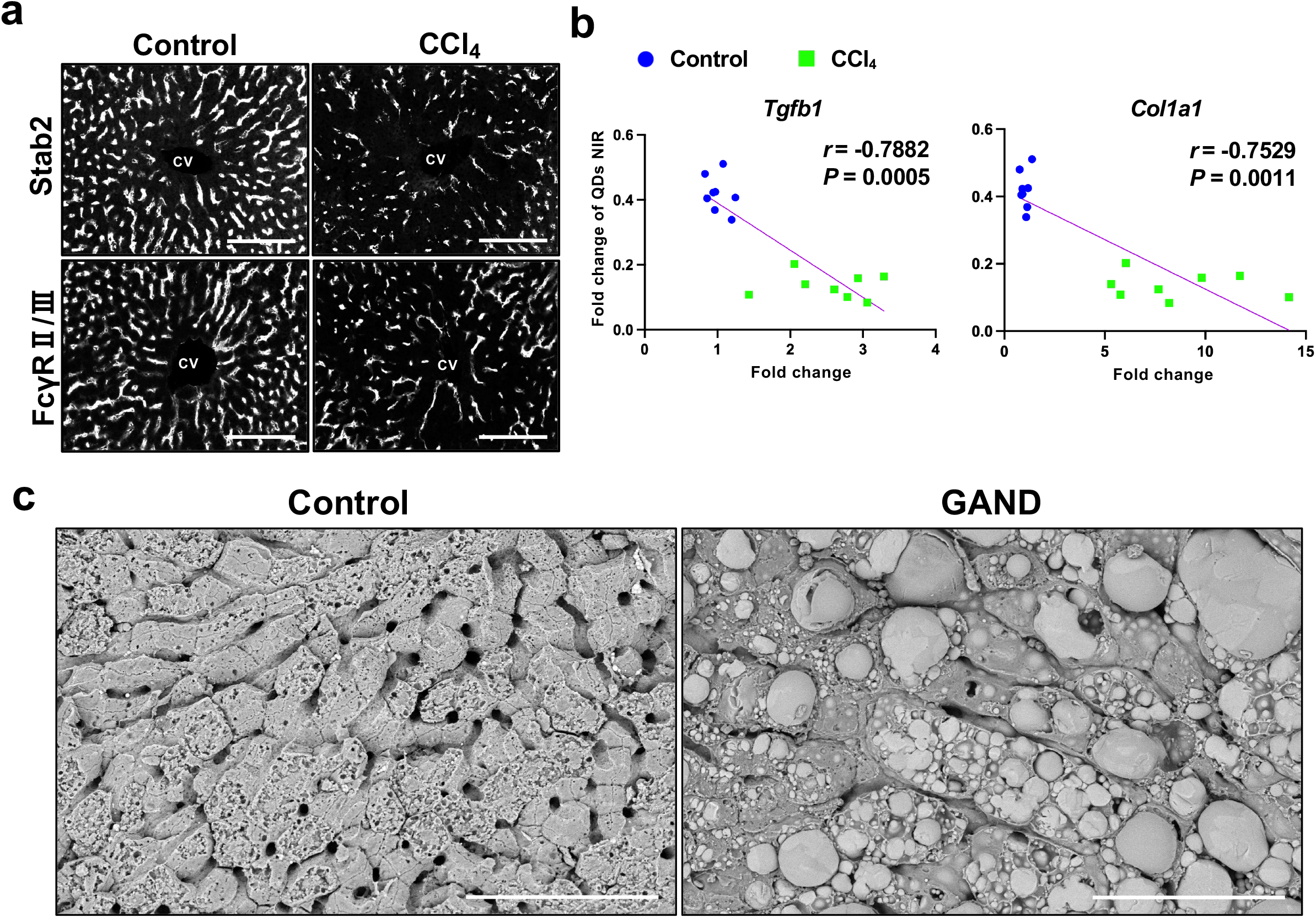
Sinusoidal injury in CCl_4_-induced acute hepatitis and GAND-induced MASH model mice. **a**, Immunohistochemical staining of Stab2 and CD32b in Control and CCl_4_-treated mouse liver sections. Scale bars, 100 μm. **b**, Scatter plots with linear trendline depicting the general relations between fold change of ǪDs NIR and pro-fibrosis marker genes (*Tgfb1* and *Col1a1*). *n* = 8 (Control and CCl_4_); *r* and *P* values were determined by spearman’s rank correlation coefficient. **c**, Scanning electron microscope (SEM) observation of liver sections extracted from Control and GAND-fed mice. Scale bars, 80 μm.

**Extended Data Fig. 3:**
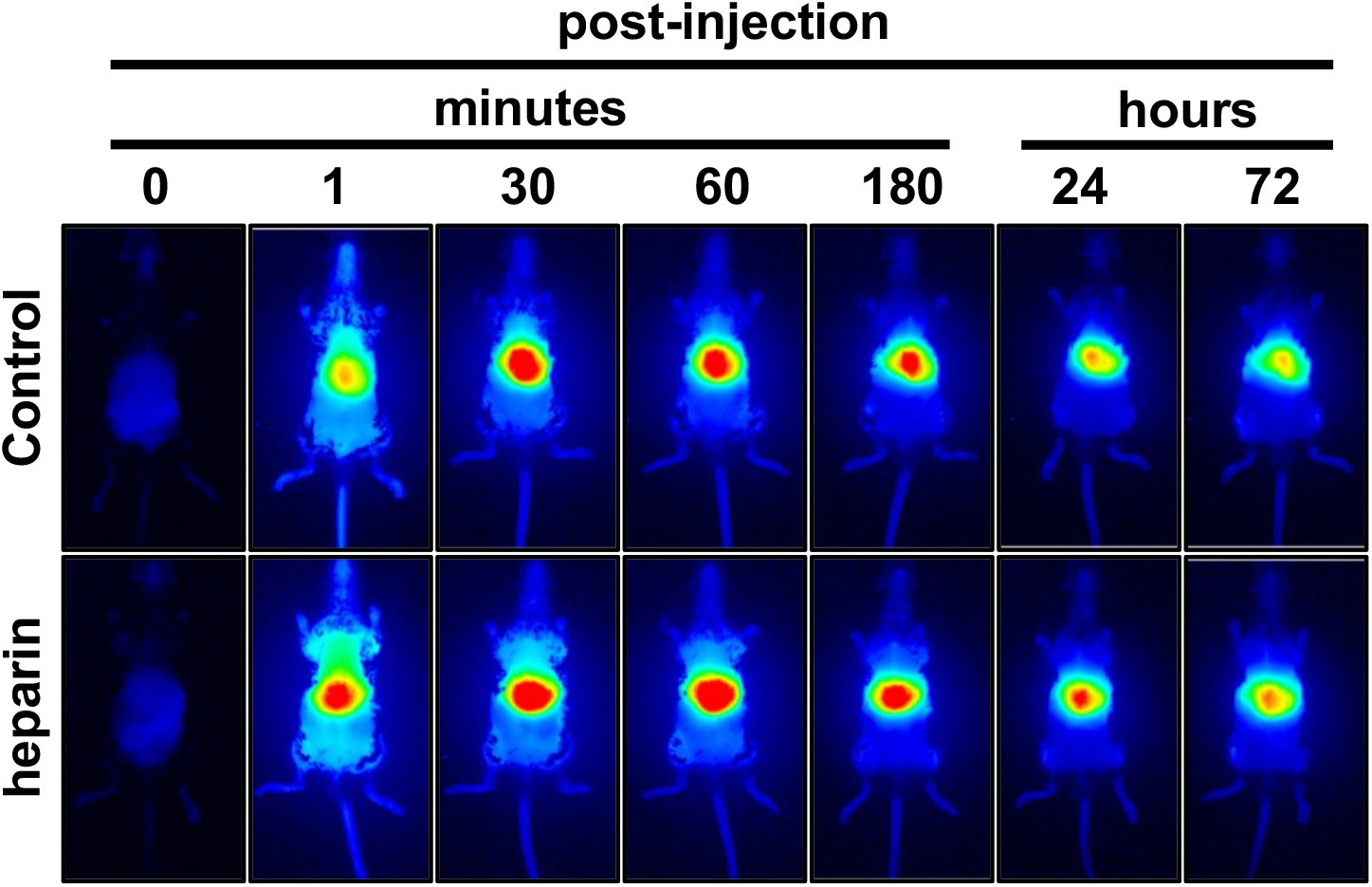
Time-lapse alteration of ǪDs NIR fluorescence detected in control and heparin-treated mice, related to Fig. 5.

**Extended Data Fig. 4:**
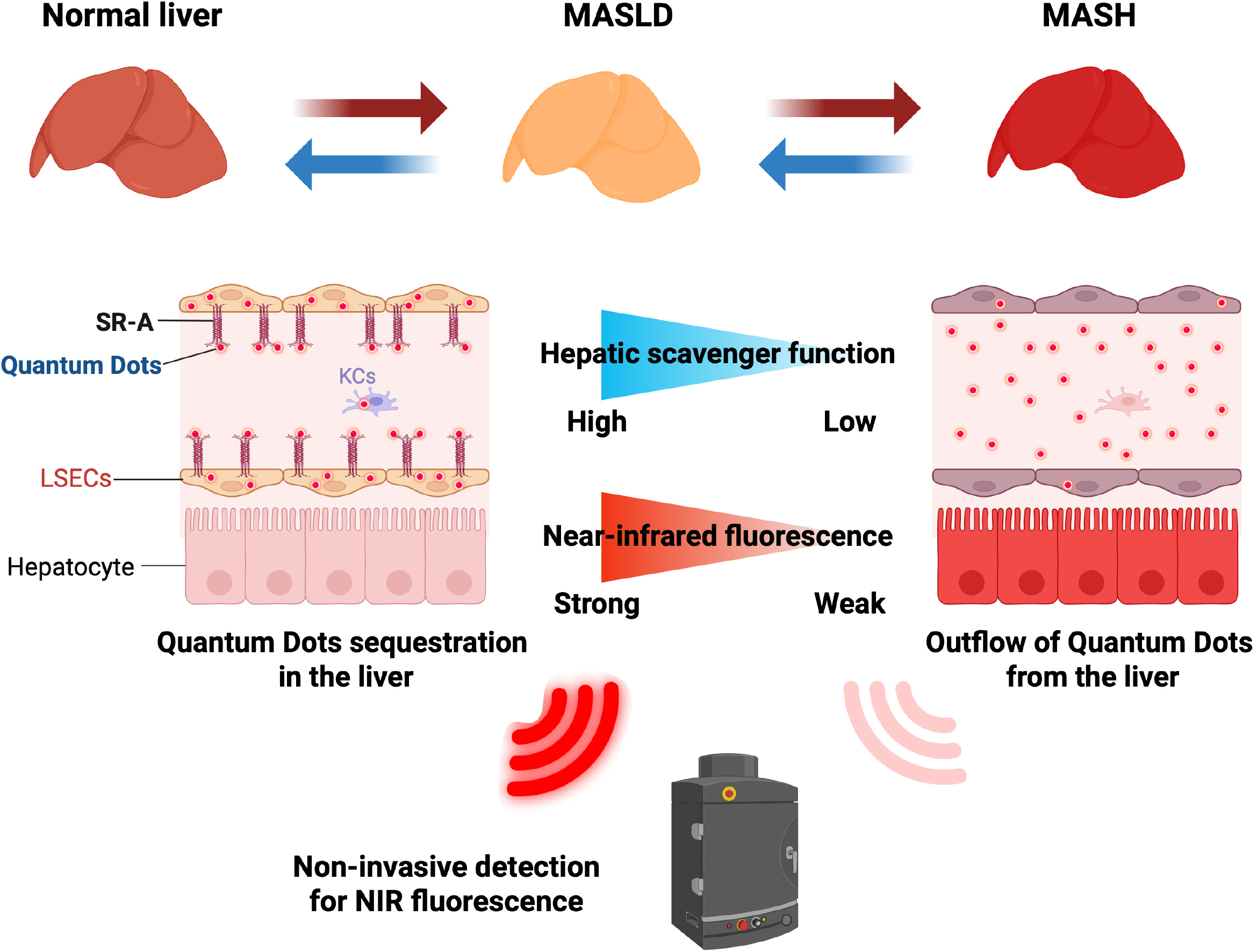
The concept of ǪDs-based non-invasive monitoring of MASH progression. Created in BioRender. Yagai, T. (2026)

## Notes

### Competing Interest Statement

The authors have declared no competing interest.

