## Supplementary Table S1 for "Quantum Dot-Based Non-invasive Monitoring of Sinusoidal Endothelial Scavenger Function in Steatohepatitis"

| Primers for mouse Quantitative RT-PCR | | |
|---|---|---|
| | Forward (5' to 3') | Reverse (5' to 3') |
| *Tnfa* | ACCCTCACACTCAGATCATCTTC | TGGTGGTTTGCTACGACGT |
| *Adgre1* | CTTTGGCTATGGGCTTCCAGTC | GCAAGGAGGACAGAGTTTATCGTG |
| *Tgfb1* | CCTGAGTGGCTGTCTTTTGACG | AGTGAGCGCTGAATCGAAAGC |
| *Col1a1* | GACATGTTCAGCTTTGTGGAC | CTTAGGCCATTGTGTATGCAG |
| *Col3a1* | TAGAGGATGGCTGTACTAAACACACTG | CAAAGACTGTCTTGCTCCATTCC |
| *Il1b* | CTGGTGTGTGACGTTCCCATTA | CCGACAGCACGAGGCTTT |
| *Gapdh* | AGGTCGGTGTGAACGGATTTG | TGTAGACCATGTAGTTGAGGTCA |
